# Interpreting differentiation landscapes in the light of long-term linked selection

**DOI:** 10.1101/131243

**Authors:** Reto Burri

## Abstract

Identifying genomic regions underlying adaptation in extant lineages is key to understand the trajectories along which biodiversity evolves. However, this task is complicated by evolutionary processes that obscure and mimic footprints of positive selection. Particularly the long-term effects of linked selection remain underappreciated and difficult to account for. Based on patterns emerging from recent research on the evolution of differentiation across the speciation continuum, I illustrate how long-term linked selection affects the distribution of differentiation along genomes. I then argue that a comparative population genomics framework that exploits emergent features of long-term linked selection can help overcome shortcomings of traditional genome scans for adaptive evolution, but needs to account for the temporal dynamics of differentiation landscapes.

---

Since the dawn of high-throughput sequencing, the inference of genomic regions of accentuated differentiation (**Box 1**) – variously referred to as differentiation islands, divergence islands, or speciations islands (**Box 2**) – has become a central focus of research on local adaptation and speciation. The past years have seen an unprecedented quest for such regions (Haasl & Payseur 2016) that assumed accentuated differentiation to evolve trough processes related to adaptation or speciation, in particular positive selection of beneficial variants (Maynard Smith & Haigh 1974; Kaplan *et al.* 1989) or selection against gene flow (Turner *et al.* 2005) in extant populations or species (in the following referred to as ‘extant lineages‘, see Glossary). However, recent research highlights that accentuated differentiation may evolve through processes other than positive selection (e. g. Bank *et al.* 2014; Cruickshank & Hahn 2014; Haasl & Payseur 2016).

### Box 1. Measures of differentiation and divergence

The interchangeable use of differentiation (allele frequency divergence) and divergence (sequence divergence) has led to their distinction as ‘relative divergence’ and ‘absolute divergence’, respectively. This distinction is important, because they describe different properties of genetic variation and are affected differently by evolutionary processes (e.g. Noor & Bennett 2009; Cruickshank & Hahn 2014). Here, for reasons of simplicity I use the former, original definition.

#### F_ST_

A measure of *differentiation* deeply rooted in population genetics theory (Whitlock 2011). It measures the relative contribution of genetic diversity between lineages to the total observed genetic diversity and is therefore dependent on genetic diversity within lineages (Charlesworth 1998). A particularly simple definition was given by Hudson et al. (1992):

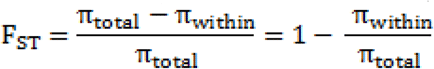

Where π_total_ and π_within_ are total nucleotide diversity and nucleotide diversity observed within lineages, respectively. In practice, F_ST_ is most often estimated using the unbiased estimator of Weir and Cockerham (1984). See Bhatia *et al*. (2013) for recommendations regarding F_ST_ estimation. It is worth noting that the π/d_XY_ ratio that has recently found application to interpret differentiation patterns (Irwin *et al.* 2016; Van Doren *et al.* 2017) is directly related to F_ST_, as π_total_ is strongly related to d_XY_.

#### d_XY_ (Nei & Li 1979)

A measure of *divergence*. It provides the average number of differences per site observed between two random haplotypes drawn from two different lineages and (in the absence of selection) is defined as: dXY = 4N_e(ANC)_μ + 2μT, where N_e(ANC)_ represents ancestral N_e_, μ the mutation rate (and thus 4N_e(ANC)_μ genetic diversity in the ancestral lineage), and T divergence time since lineage split. In practice, d_XY_ is estimated as:

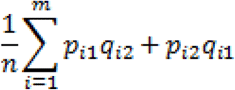

where p_i1_ and q_i1_ are the frequencies of alternative alleles at locus i in lineage 1; p_i2_ and q_i2_ the frequencies of alternative alleles at the same locus in lineage 2; m the number of variable sites; and n the sum of variable and invariable sites (importantly, following the same filtering criteria) within the interval for which d_XY_ is estimated. Contrary to F_ST_, d_XY_ is not sensitive to selection in extant lineages, but sensitive to selection in the ancestor (Nachman & Payseur 2012). At early stages of differentiation (T∼0), d_XY_ reflects ancestral diversity, 4N_e(ANC)_μ. As the contribution of 4N_e(ANC)_μ relative to 2μT diminishes, d_XY_ converges towards d (see below).

#### Relative node depth, RND (Feder *et al.* 2005)

RND is a measure of *divergence* based on d_XY_ that aims at correcting d_XY_ for mutation rate variation along the genome. To this end, it divides d_XY_ of the focal comparison by the average d_XY_ observed between the focal species and an outgroup. However, by doing so also variation in ancestral population size along the genome is corrected for. This can be problematic in species in which the recombination landscape is conserved between outgroup and focal taxa, for instance if RND is used instead of d_XY_ in F_ST_-d_XY_ contrasts.

#### d_A_ (Nei & Li 1979)

A measure of *differentiation.* This measure has been devised to capture pairwise differences that arose since the lineage split and is estimated/defined as:

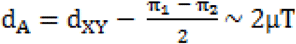

where π_1_ and π_2_ are genetic diversities observed in extant lineages 1 and 2, respectively. It has often been mistaken as a measure of divergence (see Cruickshank & Hahn 2014). Owing to the approximation of 4N_e(ANC)_μ through average diversity in descendant populations, it is dependent on genetic diversity.

#### d_f_

A measure of *differentiation*. It represents the number of fixed differences per site (first referred to as d_f_ by Ellegren et al. 2012), and therefore is readily mistaken as a measure of divergence (see Cruickshank & Hahn 2014). The evolution of the density of fixed differences is complex (Hey 1991). At early stages of differentiation, the number of fixed sites depends on the fixation of ancestral variants exclusively. d_f_ is therefore highly dependent on genetic diversity within lineages. On the long run, as fixed differences between lineages predominantly get to represent mutations which arose since lineage split, d_f_ converges towards d.

#### d (Nei 1972)

A measure of *divergence* used in the field of molecular evolution that assumes all variation between lineages to be fixed variants arisen by mutation since lineages split. As such, it is defined as d = 2μT. The estimation of d is usually based on the pairwise divergence between single sequences from two species, assuming that all variation found between sequences is fixed between species.

### Box 2. Types of genomic islands

Descriptions of genomic landscapes, in particular genomic regions of accentuated differentiation, include a number of terms that have been used interchangeably. Discussions on the existence of genomic islands (Pennisi 2014) appear of limited use. Rather, their interpretation requires a comprehensive understanding of the processes involved in the evolution of heterogeneous genomic landscapes, which is assisted by clear-cut terminology. The island metaphor has its limits in capturing the processes underlying the evolution of genomic landscapes, and interpretations of the latter may require more explicit terminology clearly distinguishing between pattern, process, and interpretation in terms of adaptation and speciation models.

#### Differentiation islands

Genomic islands exhibiting accentuated genetic differentiation (e.g. F_ST_, **Box 1**). Differentiation is sensitive to any process affecting allele frequencies, and differentiation islands can therefore evolve as a consequence of positive selection, gene flow in surrounding regions, or background selection. This term therefore has the broadest application and should be preferably used in the absence of evidence for a particular process underlying the evolution of genomic islands.

#### Divergence islands

Genomic islands exhibiting accentuated sequence divergence (d_XY_, **Box 1**). Divergence is mostly sensitive to selection in ancestral (but importantly not extant) lineages, and gene flow (for useful illustrations see (Nachman & Payseur 2012; Cruickshank & Hahn 2014)). Both processes reduce divergence, and divergence islands are therefore usually related to gene flow in the surrounding genomic regions.

#### Speciation islands

Defined as genomic regions that “remain differentiated despite considerable gene flow” (Turner *et al.* 2005). Therefore, contrary to the previous terms, relating not only to pattern, but to the supposed role in speciation. According to Nachman and Payseur (2012) and Cruickshank and Hahn (2014), speciation islands need to qualify as both differentiation islands and divergence islands. The term has found broader application such as to refer to any genomic islands involved in speciation, including genomic regions under divergent selection in context of ecological speciation in the presence of gene flow. In such cases, the usefulness of the above criterion may be limited because at these timescales divergence is determined uniquely by ancestral polymorphism (**Box 1**) (see also Burri in press). A terminology making a clearer distinction may therefore be useful. In reference to their relation to primary and secondary speciation-with-gene flow (Cruickshank & Hahn 2014), I suggest the terms ‘primary speciation island’ and ‘secondary speciation island’.

#### Incidental islands

A term coined by Turner and Hahn (2010) to describe genomic islands unrelated to speciation that evolve as a consequence of linked selection following speciation.

Particularly, several aspects of linked selection may remain underappreciated. First, awareness that purifying selection at linked sites (background selection, BGS) (Charlesworth *et al.* 1993; Charlesworth 2013) makes important contributions to linked selection and can mimic the footprints of positive selection (Stephan 2010) is still limited. Furthermore, part of the effects of linked selection observed today may have accumulated over extended periods of time and therefore be related to adaptation in ancestral rather than extant lineages (McVicker *et al.* 2009; Munch *et al.* 2016; Phung *et al.* 2016). Approaches that assist disentangling the effects of alternative forms of selection and the timescales at which they acted are called for.

Here, I first showcase the impact of linked selection on the long-term evolution of genetic diversity and hence differentiation (Charlesworth 1998). I then discuss how these effects lead to the evolution of temporally dynamic correlations of differentiation landscapes, and how this process may be influenced by demography and the evolution of genome features. I close by outlining how emergent features of long-term linked selection can be exploited to empirically take into account the long-term effects of linked selection in a comparative population genomics framework that takes into account the temporal dynamics of differentiation landscapes.

## Effects of linked selection in heterogeneous recombination landscapes

The association of physically linked genetic variants within chromosomes, and recombination – the force that can break it up – have a profound effect on the distribution of genetic diversity along genomes (Cutter & Payseur 2013). Genetic diversity becomes not only reduced at sites targeted by positive selection but also at surrounding sites. During selective sweeps, genetic variants linked to beneficial mutations hitchhike along, while others are lost (Maynard Smith & Haigh 1974). Likewise, selection against gene flow limits the levels of genetic diversity in genomic regions involved in reproductive isolation compared to introgressed proportions of the genome (Turner *et al.* 2005). Importantly, in a similar way a reduction of genetic diversity at linked sites is caused by selection against deleterious variants, i.e. BGS, although at lower rates (Charlesworth *et al.* 1993; Zeng & Charlesworth 2011). The extent to which these processes affect particular genomic regions, besides selection strength, are determined by local recombination rates (Maynard Smith & Haigh 1974; Kaplan *et al.* 1989). Specifically, in genomic regions with infrequent recombination linked selection affects physically larger chromosome stretches, and it does so at a higher rate in regions with a high density of functional sites (hereafter ‘functional density’) that constitute targets for selection (Payseur & Nachman 2002). Therefore, variation in recombination rate and in functional densities along the genome are bound to result in a heterogeneous genomic landscape of diversity, with regions of low recombination exhibiting troughs in diversity (and therefore reduced local effective population size, N_e_, as diversity π=4N_e_μ). Because differentiation is intrinsically affected by diversity (Charlesworth 1998), genomic regions of low recombination inevitably evolve towards accentuated differentiation under any of these forms of selection.

## Evolution of correlated differentiation landscapes

Recent research contributes to the emergence of a striking picture of how differentiation landscapes evolved among closely related species. *Helianthus* sunflowers, *Heliconius* butterflies, *Ficedula* flycatchers, crows, greenish warblers (*Phylloscopus trochiloides*), and stonechats (**Fig. 1**) exhibit highly similar differentiation landscapes among closely related species (Kronforst *et al.* 2013; Martin *et al.* 2013; Renaut *et al.* 2014; Burri *et al.* 2015; Irwin *et al.* 2016; Vijay *et al.* 2016; Van Doren *et al.* 2017).

**Figure 1.**
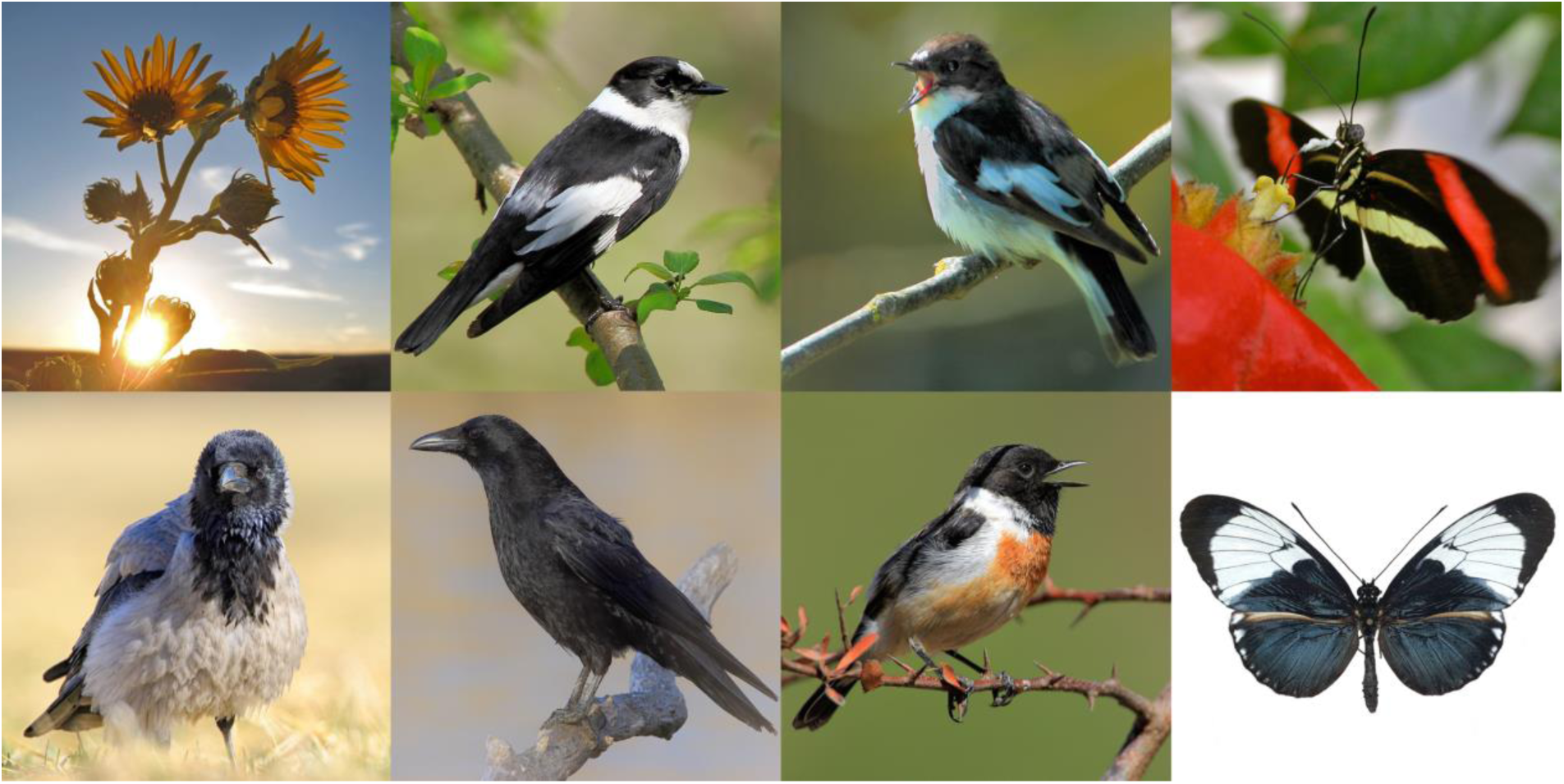
Systems with correlated differentiation landscapes. From top left to bottom right: *Helianthus* sunflowers, collared flycatcher (*Ficedula albicollis*), pied flycatcher (*Ficedula hypoleuca*), *Heliconius melpomene rosina*, hooded crow (*Corvus cornix*), carrion crow (*Corvus corone*), Siberian stonechat (*Saxicola maurus*), *Heliconius cydno galanthus* (picture credits: sunflowers, Takeshi Kawakami; *Heliconius* butterflies, Richard Merrill; birds, Reto Burri)

This pattern of repeated evolution is a direct prediction of the long-term action of linked selection in heterogeneous recombination landscapes. Over timescales across which the genome features constraining genetic diversity are stable, diversity landscapes are expected to correlate even among independent lineages for two reasons. First, diversity levels along the genome scale with levels of ancestral diversity (**Fig. 2A-B**). This cumulative impact of linked selection is reflected in a correlation of extant diversity (*π*) with ancestral diversity (reflected in sequence divergence, *d*_*XY*_, among closely related species, **Box 1**) (Nachman & Payseur 2012; Burri *et al.* 2015). The reduction of ancestral diversity through long-term linked selection may be strong enough to even impact sequence divergence between distantly related species, such as human and mouse, and bird species as divergent as 50 my (Phung *et al.* 2016; Dutoit *et al.* 2017; Van Doren *et al.* 2017; Vijay *et al.* 2017). Second, linked selection within extant lineages maintains the relative levels of diversity among genomic regions. This is reflected most directly in reduced levels of private polymorphisms in low-recombination regions, as observed in flycatchers (Burri *et al.* 2015). Importantly, despite extant diversity in part being explained by ancestral diversity, correlated *differentiation* landscapes can evolve independently; differentiation evolves only through the differential sorting of genetic diversity in independent extant lineages. In conclusion, the evolution of correlated differentiation landscapes is an effect of long-term linked selection exposing conserved genomic features, namely conserved landscapes of recombination and functional densities.

**Figure 2.**
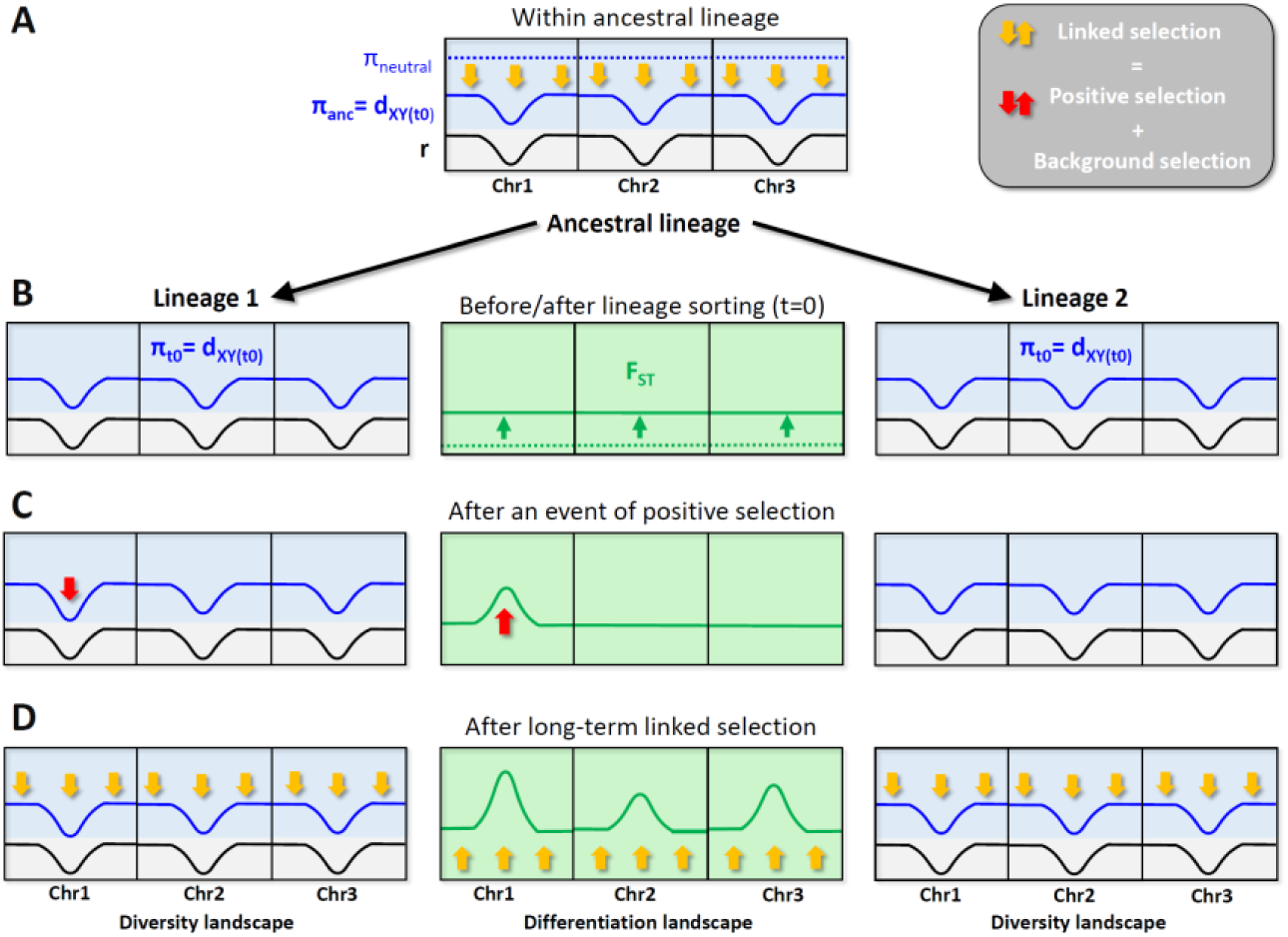
Genetic diversity (π, blue shaded boxes) and differentiation (F_ST_, green shaded boxes) from early to late stages of differentiation and the role of linked selection in their evolution. Three example chromosomes are depicted. The grey shaded parts of the boxes represent the underlying recombination landscapes. **A)** Genetic diversity in a hypothetical population with a heterogeneous recombination landscape before (broken line) and after (solid line) the action of any form of selection. **B)** Genetic diversity and differentiation after a population split. Differentiation is shown before (broken line) and after lineage sorting (solid line) with lineage sorting depicted by green arrows. Before lineage sorting, genetic diversity in the descendent populations (π_t0_) equals ancestral diversity (d_XY(t0)_ = 4N_e(ANC)_μ + 2μT = 4N_e(ANC)_ μ). After some amount of lineage sorting, differentiation starts building up genome-wide. At this timescale, background selection had limited power to reduce diversity and enhance differentiation in low-recombination regions, and no event of positive selection may have occurred. **C)** After the occurrence of an event of positive selection in one population, differentiation is enhanced in one chromosome. At this timescale, background selection still had limited power to reduce diversity and enhance differentiation in low-recombination regions. **D)** As a long-term effect of linked selection (including both positive selection and BGS) differentiation starts building up more strongly in all low-recombination regions than in the remainder of the genome.

Given that BGS explains a high proportion of the variation in diversity along genomes (Lohmueller *et al.* 2011; Comeron 2014; Corbett-Detig *et al.* 2015; Phung *et al.* 2016), in finite populations that are mutation-limited, BGS may arguably play a major role in the evolution of these patterns. Beneficial mutations are rare (Eyre-Walker & Keightley 2007), and the frequency at which positive selection hits the same genomic regions repeatedly is expected to be limited. In contrast, deleterious variants represent the majority of non-neutral mutations. With a steady influx of deleterious mutations, the cumulative time span during which BGS acts is expected to be much longer than that of episodic positive selection, and mutations representing targets for BGS are abundant and widespread across the genome. Consequently, whilst being pervasively affected by BGS, not all functional regions are necessarily hit by events of positive selection. Therefore, *in extremis*, genomic landscapes shared among species may evolve even under the action of BGS alone.

Even though beneficial mutations are rare, at vast timescales there is an increasing chance for several of them to occur within the same genomic region. In such cases, also positive selection may affect the same genomic regions repeatedly. However, it may often have occurred at timescales exceeding lineages’ divergence times, and therefore involve events of positive selection other than ones associated with adaptive evolution in extant lineages.

It appears that on the long run low-recombination regions are predestined to evolve towards accentuated differentiation due to the cumulative effects of pervasive BGS and, presumably to a lesser extent, recurring sweeps (McVicker *et al.* 2009; Munch *et al.* 2016). As a consequence of this process, conservation of recombination rates and functional densities among lineages will lead to the evolution of correlated diversity and differentiation landscapes even among distantly related lineages.

## Unpredictable emergence of differentiation islands at early stages of differentiation

The existence of genomic regions predestined to evolve accentuated differentiation prompts the question whether the same patterns are already present at early stages of differentiation. Indeed, sharing of highly differentiated regions at early stages of differentiation has been reported from diverse organisms (Jones *et al.* 2012; Renaut *et al.* 2013; Soria-Carrasco *et al.* 2014; Fraser *et al.* 2015). Moreover, in the few studies that investigated multiple timescales, regions shared among older comparisons are on average also more differentiated earlier on. In flycatchers and crows (Burri *et al.* 2015; Vijay *et al.* 2016), regions exhibiting accentuated differentiation among populations do so also between species. However, vice versa, only a subset of regions exhibiting accentuated differentiation between species do so among populations. Similar patterns are found in *Heliconius* butterflies (Kronforst *et al.* 2013; Martin *et al.* 2013), and in *Helianthus* sunflowers (Renaut *et al.* 2014). These observations suggest that the effects of linked selection expose low-recombination regions less homogeneously at early stages of differentiation than in the long run.

Current evidence suggests that at early stages of differentiation BGS has limited power to explain accentuated differentiation. The effects of BGS may be too subtle to drive accentuated differentiation at short timescales. Otherwise, differentiation would be predicted to evolve in identical genomic regions in related lineages (Munch *et al.* 2016). However, this expectation is not met by observations from early differentiation stages in flycatchers and crows: genomic regions highly differentiated among populations within one flycatcher species or across one crow hybrid zone usually exhibit average differentiation in other species/hybrid zones (Burri *et al.* 2015; Vijay *et al.* 2016). The emergence of differentiation islands (**Box 2**) at these rather recent timescales, therefore, appears to be unpredictable. In particular the observation that the locations of such regions differ between closely related species in flycatchers and across crow hybrid zones suggest that, apart from the effects of drift connected with lineage splits, the evolution of accentuated differentiation at early stages of differentiation may involve positive selection rather than BGS.

## A temporal perspective: Dynamic correlations among genomic landscapes along the differentiation continuum

The outlined observations suggest temporal dynamics of differentiation landscapes during which (i) differentiation increasingly reflects underlying recombination rates and functional densities, and correlations among independent differentiation landscapes increase with advancing differentiation, and (ii) the interpretation of accentuated differentiation may depend on the differentiation stage in focus. These temporal dynamics have important bearings on the design and interpretation of population genomic studies.

In populations evolving under neutrality, genetic diversity (π) is unaffected by recombination rate variation (**Fig. 2A**). However, selection reduces genetic diversity, with strongest effects on low-recombination regions. The resulting heterogeneous diversity landscape is passed on to descendant lineages. Shortly after the lineages split, differentiation will be minimal, mainly driven by the sampling effect connected with the split (**Fig. 2B**). In a first stage, differentiation will then start building up genome-wide, predominantly through differential lineage sorting of ancestral polymorphisms. At these early stages of differentiation, BGS has made limited contributions to enhance lineage sorting in low recombination regions. Therefore, unless the effects of genetic drift connected with lineage splits are unusually pronounced in particular genome regions (see next section), the evolution of accentuated differentiation may require positive selection (**Fig. 2C**). As a function of the waiting times for positive selection to occur, the emergence of regions of strongly accentuated differentiation may be unpredictable, and the correlation of differentiation landscapes among independent comparisons remain limited (**Fig. 3A**).

**Figure 3.**
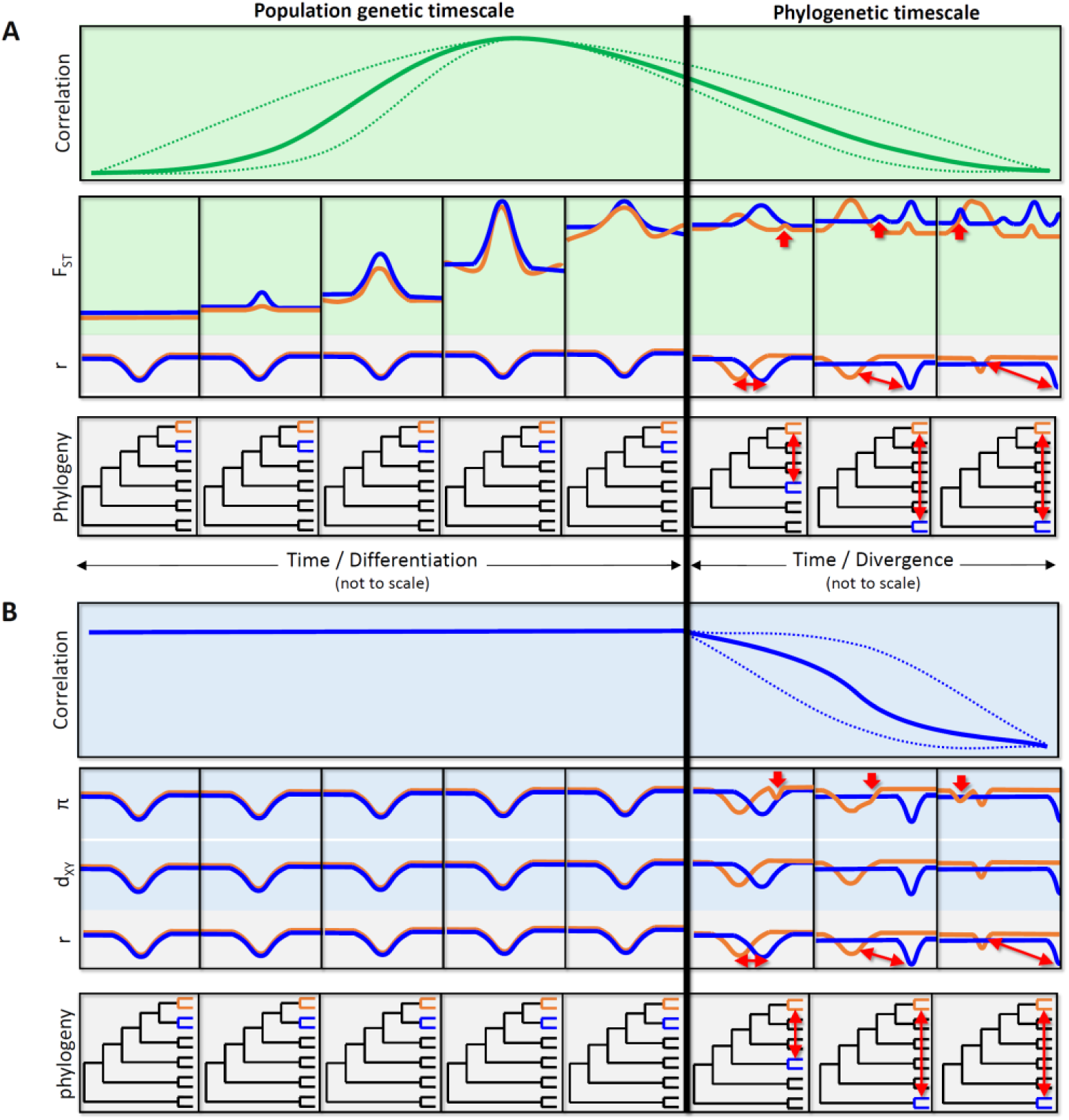
Sketch of temporal dynamics of the correlation among independent differentiation, diversity, and divergence landscapes with increasing differentiation and divergence. **A**) Temporal dynamics of differentiation landscapes and their correlation. Top panel: Temporal dynamics of the correlation of differentiation landscapes. The form of this correlation is not clear, as indicated by the alternative trajectories (broken lines). Middle panel: Evolution of the underlying differentiation and recombination landscapes of the two comparisons (depicted in blue and orange; blue Comparison 1; orange, Comparison 2). Bottom panel: Phylogenetic topologies indicating the divergence between the two pairwise comparisons. **B**) Temporal dynamics of the genomic landscapes of diversity (π) and sequence divergence (d_XY_). The temporal dynamics of the correlations in π and d_XY_ among lineages/comparisons are expected to be similar, for which reason only one temporal dynamics is shown (upper panel). π and d_XY_ are not affected the same way as F_ST_ by lineage sorting after a lineage split and therefore are expected to be correlated throughout the population genetic timescale. Note that selection in extant lineages does not affect d_XY_. In both panels the temporal dynamics at the population genetic (left) and phylogenetic timescales (right) are shown. Axes are not to scale. Fat red arrows, lineage-specific sweeps; slim red arrows, divergent evolution of the recombination landscapes, and divergence between comparisons.

As differentiation advances, the effects of linked selection will start accumulating. and differentiation landscapes will increasingly reflect the underlying recombination landscape, such as illustrated, e.g., in stickleback fish (Roesti *et al.* 2013). Compared to earlier stages of differentiation, these long-term effects of linked selection will contribute to a fundamentally different picture characterized by consistently accentuated differentiation in low-recombination regions (**Fig. 2D**). Across time scales for which the recombination landscape is stable – this can be across numerous speciation events, and tens of millions of years in birds (Singhal *et al.* 2015) – this process will result in strong correlations among differentiation landscapes from independent comparisons (**Fig. 3A**). In contrast to shorter timescales, the cumulative effects of BGS had more time to play out, and accentuated differentiation may evolve even in the absence of positive selection.

Altogether, this leads to temporally dynamic correlations among differentiation landscapes from independent comparisons (**Fig. 3A**). In contrast, at population genetic timescales (**Fig. 3** left) no such temporal dynamics are expected for the genomic landscapes genetic of diversity (π) and sequence divergence (d_XY_), because these are passed down over lineage splits (**Fig. 3B**).

However, in the long run, the correlations among genomic landscapes are eventually bound to decay along two axes: (i) a population genetic timescale, i.e. the stage of differentiation *within* each pairwise comparison (**Fig. 3** left), and (ii) a phylogenetic timescale, i.e. the phylogenetic divergence *between* pairwise comparisons (**Fig. 3** right). Within the first, the variance in differentiation (F_ST_, **Box 1**) is reduced as it reaches unity towards the end stages of differentiation. Stochastic variation may therefore lead to decreasing correlations at this stage (**Fig. 3** right). Perhaps more importantly, at the phylogenetic timescale, as divergence time between the differentiation landscape increases, even relatively stable recombination landscapes are bound to diverge, and lineage-specific events of positive selection accumulate. Both processes will contribute to erode the correlation of differentiation landscapes with increasing divergence (**Fig. 3**). The divergent evolution of recombination will also contribute to erode the correlation among independent landscapes of diversity and sequence divergence (**Fig. 3B**) over phylogenetic timescales.

## The complex interplay of genomic features and population-level processes in the evolution of differentiation landscapes

The outlined temporal dynamics may vary strongly, and depend on the complex interplay of various genomic features and population-level processes.

Recombination rate evolution differs substantially among species (Smukowski & Noor 2011), and may have a strong impact on the evolution of genomic landscapes. First, among taxa in which the amplitude of recombination rate variation is limited, random variation in genetic diversity and differentiation may dominate and differentiation landscapes may correlate poorly (**Tab. 1**). Second, the conservation of recombination landscapes, determines the timescale across which the same (low-recombination) regions are exposed to strong effects of linked selection. In taxa with long-term conserved recombination landscapes, such as birds (Singhal *et al.* 2015), diversity reductions through linked selection in low-recombination regions will be particularly strong, and diversity and differentiation landscapes will be correlated over longer timescales than in taxa with more dynamically evolving recombination landscapes, such as most mammals (e.g. Oliver *et al.* 2009; Baudat *et al.* 2010) (**Tab. 1**).

How other genomic features affect the evolution of correlated differentiation landscapes may depend on the extent and sign of their correlation with recombination rate and the turnover of their distribution along the genome (**Tab. 1**). Mutation rate can vary substantially (e.g. Hodgkinson & Eyre-Walker 2011) and appears to be elevated in regions of high recombination (e.g Cutter & Payseur 2013; Arbeithuber *et al.* 2015; Terekhanova *et al.* 2017). Although such a positive correlation of mutation with recombination might reinforce the effects of linked selection, there is little evidence for such a concerted effect on genetic diversity (Francioli *et al.* 2015; Ellegren & Galtier 2016). Meanwhile, functional densities are positively correlated with recombination rate to varying degrees (e.g. Flowers *et al.* 2012; Nam & Ellegren 2012; Kawakami *et al.* 2014). In taxa in which this antagonistic correlation is strong, it may dampen the effects of linked selection (Cutter & Payseur 2013). Moreover, a quick turnover of hot- and cold-spots of mutation and functional densities would distribute the respective effects more homogeneously along the genome in the long run. The turnover of functional densities may be moderate; even though new genes emerge at high rates, the majority is short-lived (Schlötterer 2015). Meanwhile mutation rates appear to evolve at a faster pace than recombination (Terekhanova *et al.* 2017), and may therefore have more ephemeral effects on the heterogeneity of genomic landscapes than long-term linked selection in conserved recombination landscapes.

In addition to genome features, population-level processes and parameters, in particular demography and genome-wide N_e_, may play a crucial role in shaping differentiation landscapes (**Tab. 1**). Even though demographic effects are expected to reduce genetic diversity genome-wide, they may have particularly strong effects on low-recombination regions. First, the higher linkage among sites results in an elevated variance of diversity and differentiation even under neutral evolution. Even though at early differentiation stages, this may result in accentuated differentiation to be mistaken as a footprint of positive selection, by itself it does not increase differentiation in low-recombination regions systematically. Second, however, low-recombination regions affected by long-term linked selection are characterized by reduced N_e_. Demographic effects may therefore contribute to systematic increases in differentiation in these regions, and reduce the correlation between differentiation landscapes of taxa with unequal demographic histories.

Finally, correlated genomic landscapes may only evolve in taxa with sufficiently high N_e_, in which selection is not overwhelmed by genetic drift (Ohta 2002) (**Tab. 1**). In line with this, linked selection appears to reduce genetic diversity below neutral expectations across a wider range of recombination rates in taxa with high N_e_ (Corbett-Detig *et al.* 2015). In taxa with very low N_e_, effects of linked selection may be negligible altogether, and correlations between genomic landscapes be absent, or even inversed (Van Doren *et al.* 2017; Burri in review)

In conclusion, the alignment of multiple genomic features in space (along the genome) and over time in combination with population-level processes are expected to determine the conditions under which correlated differentiation landscapes may evolve and the timescales over which correlations persist (see **Outstanding questions**).

## Accounting for the long-term effects of linked selection: a comparative population genomics framework

While genomic regions of accentuated differentiation have commonly been interpreted as a footprint of ecological adaptation or selection against gene flow, awareness is increasing that such an approach is naïve towards the vast recombination rate variation found in many species, the long-term effects of linked selection associated therewith, and the contribution of BGS therein. Distributions against which to discriminate outliers need to be adapted, and, as suggested by the temporal dynamics of differentiation landscapes outlined above, timescales may need to be accounted for.

At early stages of differentiation, genome-wide differentiation may offer a baseline against which to discriminate candidate regions (but see below). However, later on, owing to the long-term effects of linked selection, this baseline is shifted upwards in low-recombination regions (**Fig. 2D**). With recombination rate data and functional annotations at hand, it is possible to devise comprehensive models that estimate parameters of BGS along the genome to adapt baselines accordingly and enable the discrimination of positive selection against BGS (McVicker *et al.* 2009; Elyashiv *et al.* 2016; Huber *et al.* 2016). In the absence of such data, alternative approaches are required.

In line with previous suggestions, I advocate that a comparative genomic framework may significantly assist the inference of candidate regions involved in adaptation and/or speciation (Berner & Salzburger 2015). First, this approach provides information on whether or not constraints imposed by long-term linked selection apply to the taxa of interest. Second, the correlation of genomic landscapes among independent comparisons at advanced stages of differentiation can be exploited to expose the (relative) baseline levels of diversity and differentiation expected to evolve under long-term linked selection.

In the simplest form of this approach, observations across multiple comparisons serve as *qualitative* ‘evolutionary controls’: Differentiation patterns common to all (focal and control) comparisons constitute a baseline hypothesis that assumes the evolution of common patterns through long-term linked selection. Only accentuated differentiation specific to single lineages may be taken to suggest positive selection in extant lineages (in the focal lineage or in the ancestor of the remaining lineages if they share a common ancestor to the exclusion of the focal lineage). Ideally, this criterion is applied even at early stages of differentiation to safeguard against false positives in regions of exceptionally low recombination and high functional density that may evolve accentuated differentiation most rapidly.

This qualitative criterion rigidly treats genomic regions with accentuated differentiation in all comparisons as unrelated to positive selection. However, in genomic regions strongly affected by long-term linked selection the effects of lineage-specific positive selection accumulate on top of already accentuated differentiation. The qualitative criterion may therefore be overly conservative. Instead, one might rather want to determine the varying amplitude to which baseline levels of differentiation need to be shifted along the genome. At differentiation stages at which differentiation landscapes are strongly correlated, the differentiation patterns common to independent comparisons are expected to reflect the long-term effects of linked selection (Munch *et al.* 2016), and may therefore be used as *quantitative* evolutionary controls. A dynamic baseline across the genome taking the effects of long-term linked selection into account can therefore be directly formulated based on the empirically observed common pattern. Several flavors of this approach can be envisaged (**Box 3**) and have helped to focus research on refined sets of candidate regions (e.g. Roesti *et al.* 2012; Roesti *et al.* 2013; Vijay *et al.* 2016).

#### Box 3. Empirical approaches for the inference of lineage-specific accentuated differentiation using quantitative evolutionary controls

Amongst the first who recognized that differentiation landscapes followed a discernible pattern along chromosomes, Roesti et al. (2012) accounted for generally high differentiation in chromosome centers observed among stickleback populations pairs using residual differentiation (F_ST_’) after accounting for chromosome-wide large scale differentiation by smoothing raw F_ST_. Following this approach that considered single comparisons, Berner and Salzburger (2015) suggested to account for differentiation patterns resulting from BGS in low-recombination regions by adjusting observed differentiation for background differentiation observed across multiple comparisons. This comparative approach using so-called delta differentiation (ΔF_ST_) has found application, e.g., in crow speciation genomics work (Vijay *et al.* 2016) to formulate baseline levels of differentiation based on mean differentiation estimated from a reference set of independent comparisons, and was extended to other population genetic parameters, such as linkage and haplotype structure in research on stickleback adaptation (Roesti *et al.* 2015).

This approach empirically adapts baselines across the genome to take into account heterogeneity in population genetic parameters due to processes unrelated to ecologically relevant lineage-specific positive selection. The extent to which this baseline reflects common underlying processes is expected to scale with the strength of correlation among independent observations (and by the correlation of differentiation landscapes with the underlying diversity landscapes), which is not quantified by the above approach. Principle component analysis (PCA) of multiple differentiation landscapes may offer an interesting alternative in this respect: Values along the first axis (PC1), similar to mean F_ST_, provide expected relative values of differentiation along the genome, while the proportion of variance explained by the axis provides information on the correlation among differentiation landscapes. Outlier regions can then be inferred from type II regression of observed differentiation against PC1. The pairwise nature of F_ST_ complicates this approach, and lineage-wise estimates of differentiation, such as the population branch statistic (PBS; Shriver *et al.* 2004; Yi *et al.* 2010) or population-specific F_ST_ (e.g. Buckleton *et al.* 2016), or nucleotide diversity (π) which is tightly bound to F_ST_, may be better suited for this kind of analysis.

Approaches like these are expected to yield sets of genomic regions enriched for candidate regions that evolved under positive selection. To verify this prediction, the comparison of distributions of parameters more explicitly related to positive selection, such as haplotype structure (Voight *et al.* 2006; Sabeti *et al.* 2007; Tang *et al.* 2007; Ferrer-Admetlla *et al.* 2014) among candidate regions and the genomic background is helpful (e.g. Roesti *et al.* 2015). Furthermore, parameter distributions within genomic regions with consistently accentuated differentiation across independent comparisons may constitute more conservative baselines against which to discriminate candidate regions than the genomic background.

**Figure Box 3.**
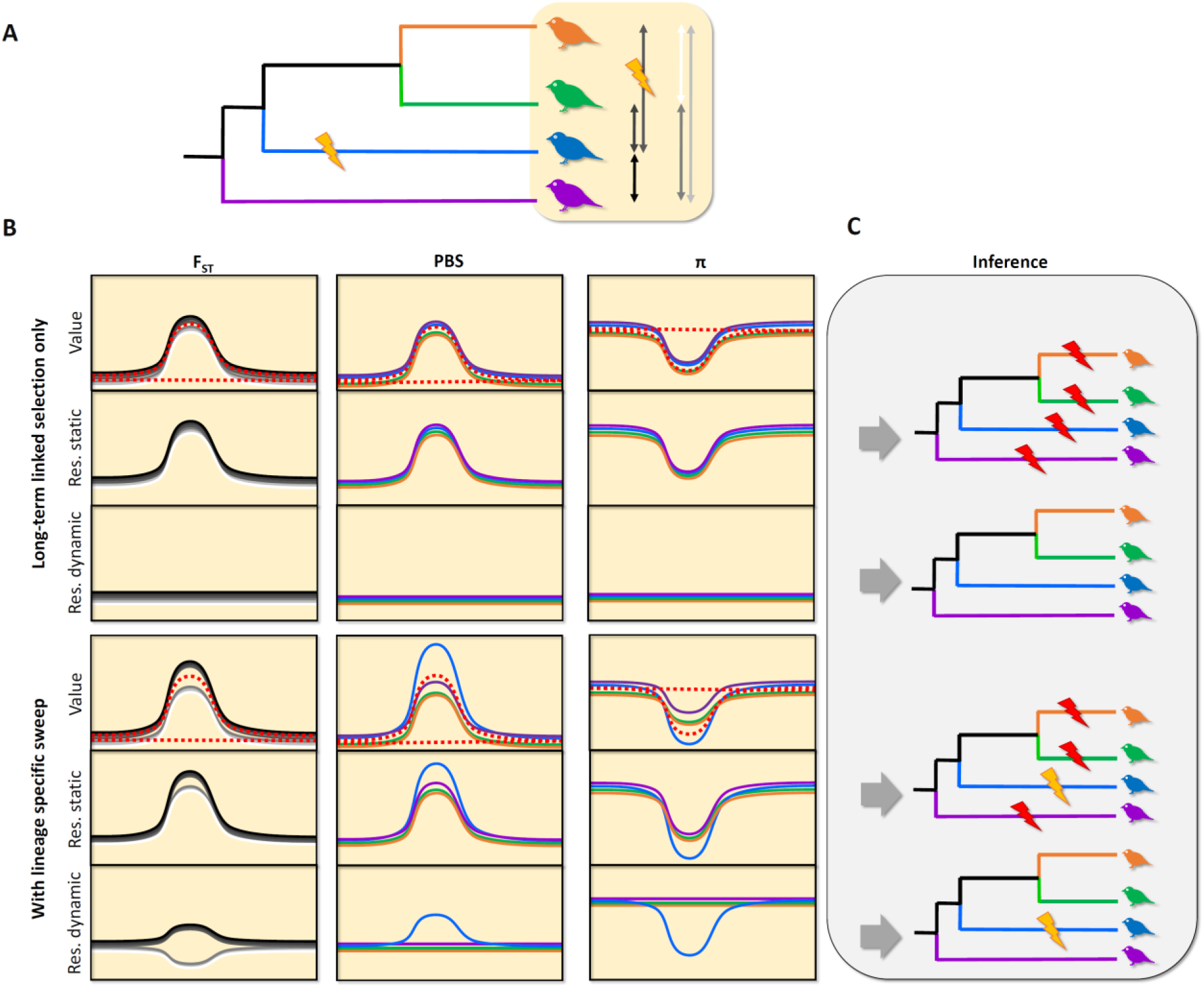
Comparative population genomics approach. **A)** Phylogenetic relationships among four species. A selective sweep is indicated on the branch leading to the blue species. Pairwise distances are indicated to the right, separated according to whether (left) or not (right) they would be affected by the indicated sweep. **B)** Pairwise-differentiation (F_ST_, with grey shades following the ones in A), lineage-specific differentiation (PBS), and diversity (π) on a chromosome with low recombination in the middle, and their residuals from a static baseline (flat broken red line) and a dynamic baseline (curved broken red line). These are provided for both a scenario without (top panels) and with (bottom panels) the selective sweep depicted in A). **C)** Inference of selective sweeps based on residuals. Red flashes indicate false positive inference of selective sweeps. A static baseline leads to elevated residuals in the chromosome center and thereby false positive inference of sweeps in all instances. With a dynamic baseline only lineage-specific elevations/reductions of differentiation/diversity are reflected as positive residuals and this inferred as footprints of a sweep. Note that with a dynamic baseline, due to its pairwise nature, F_ST_ results in a blob of negative and positive residuals in the presence of a sweep (lower left), and population-specific measures of differentiation or diversity may be better suited for this approach.

##### Outstanding Questions

To identify the footprints of adaptation and speciation in the differentiation landscape, an improved understanding of the contributions of BGS, positive selection in the ancestor, and their interaction with demography towards the evolution of differentiation landscapes and their correlation is required. Previous simulations have shown that BGS in extant populations substantially affects genetic diversity and differentiation, with strongest effects close to the sites it targets and in genomic regions where recombination is infrequent, resulting in local reductions of extant N_e_ within the genome (Zeng & Charlesworth 2011; Zeng & Corcoran 2015) that can be mistaken as a footprint of positive selection (Huber *et al.* 2016). These simulations suggest that BGS acting within populations may not reduce diversity by more than two thirds relative to neutral expectations (Zeng & Charlesworth 2011) and may thus not explain the diversity reductions observed for instance in greenish warblers (see e.g. Irwin *et al.* 2016). Moreover, the effects of BGS in extant lineages must be stronger than in the ancestor to result in elevated differentiation (Zeng & Corcoran 2015). However, recent results suggest that the effects of BGS particularly in ancestral lineages may have been underestimated so far (Phung *et al.* 2016). Moreover, the effects of demography acting upon a heterogeneous diversity landscape remain largely unexplored. With regard to the model proposed here a number of question remain open that would profit from additional simulation and empirical work:

- How strongly can BGS reduce genetic diversity in low-recombination regions relative to the remainder of the genome, if it acts over multiple subsequent speciation events over millions of years in taxa with long-term conserved recombination landscapes?
- At which timescales do correlations among differentiation landscapes build up? In particular, from which stage of differentiation onwards are effects of BGS discernible in different bins of recombination rate and functional densities, and what are the long-term contributions of BGS and recurrent hitchhiking?
- What is the effect of demographic events, such as repeated cycles of range expansions and contractions, on heterogeneous diversity landscapes produced by linked selection? Are the resulting reductions in diversity (and N_e_) overproportional in regions with already reduced N_e_, and may demographic events thus further amplify the heterogeneity of differentiation landscapes in addition to the effects of linked selection?
- How do recombination rate evolution, lineage-specific episodes of positive selection, and variation in effective population size among lineages interplay in the temporal dynamics of build-up and decay of the correlation among independent differentiation landscapes?
- How strong do demographic effects, such as bottlenecks, need to be to perturb the evolution of correlated differentiation landscapes?
- In how far can the evolutionary dynamics of differentiation landscapes be extrapolated among various organisms? Which roles may the mechanisms determining recombination rates have therein?

## Temporal dynamics of differentiation landscapes in the comparative population genomics framework

Although both qualitative and quantitative evolutionary controls have found application (e.g. Roesti *et al.* 2012; Roesti *et al.* 2013; Renaut *et al.* 2014; Feulner *et al.* 2015; Vijay *et al.* 2016), the outlined temporal dynamics open an issue that has found less attention: the choice of appropriate evolutionary controls. I argue that these need to be chosen carefully with respect to both the stage of differentiation and the phylogenetic divergence.

Mismatched evolutionary controls may result in high false discovery rates. For comparisons at early differentiation stages (**Fig. 2 B-D**, **Fig. 3** left) controls at an advanced stage (**Fig. 2D**, **Fig. 3** right) risk yielding high rates of false negatives; although in some genomic regions accentuated differentiation may be bound to evolve in the long run, at early differentiation stages it may require positive selection. Conversely, for comparisons situated at advanced differentiation stages, comparisons at earlier stages, at which the effects of BGS may not yet be discernible, are prone to yield false positives. Similar problems may be induced by exceedingly divergent evolutionary controls (**Fig. 3** right), which are prone to bias inference towards false positives. Due to an increased risk of divergent recombination landscapes, such divergent controls may not adequately reflect the long-term effects of linked selection relevant for the comparison. Consequently, evolutionary controls may yield most conclusive results when chosen at differentiation stages similar to that of comparisons, and in close phylogenetic proximity to the latter.

Nevertheless, finding adequate evolutionary controls may be complex. In many instances differentiation and divergence parameters may not be known when setting out for population genomics experiments. Ideally, experimental designs therefore may incorporate a range of evolutionary controls strategically placed along the differentiation continuum.

## Conclusions

Long-term linked selection in heterogeneous recombination landscapes leads to the evolution of highly heterogeneous differentiation landscapes. This process results in temporally dynamic correlations among independent differentiation landscapes of related lineages, within which the interpretation of accentuated differentiation in terms of positive selection in extant lineages is complicated to different extents depending on the timescale studied. The comparative population genetic approach provides a powerful tool to qualitatively or quantitatively adapt baselines along the genome to take into account the long-term effects of linked selection. However, evolutionary controls have to be matched to the stage of differentiation and chosen at appropriate phylogenetic divergence. Empirical research and simulations are now called for to investigate the temporal dynamics of correlations among independent differentiation landscapes (see **Outstanding questions**), and to study the interplay of different forms of selection, recombination rate variation, and demography in their evolution.

**Table 1.** Conditions under which genomic features and population-level processes may favour or disfavour the evolution of correlated differentiation landscapes.

|  | **Favours correlation** | **Disfavours correlation** |
| --- | --- | --- |
| **Recombination rates** | Highly heterogeneous along the genome<br>Conserved across long timescales | Homogeneous along the genome<br>Quick turnover |
| **Functional densities** | Negative correlation with recombination rates<br>Conserved across long timescales | Positive correlation with recombination rate<br>Quick turnover |
| **Mutation rates** | Positive correlation with recombination rate *and* conserved across long timescales | Negative correlation with recombination rate<br>Quick turnover |
| **N<sub>e</sub>** | High N<sub>e</sub> | Low N<sub>e</sub> |
| **Demography** | Similar demography among lineages | Lineage-specific demographic events |

## Glossary

Background selection: A process by which purifying selection against deleterious variants results in a loss of neutral genetic diversity at linked sites.
Extant lineages (extant populations, extant species): Non-ancestral lineages, i.e. lineages observed today including the time since the split from their ancestor.
Genomic landscapes: The distribution of parameters, such as diversity (‘diversity landscape’), differentiation (‘differentiation landscape’) or recombination rates (‘recombination landscape’) along chromosomes.
Hitchhiking/recurrent hitchhiking: A process by which positive selection for a beneficial variant leads to an increase in frequency of neutral variants at linked sites, and thus loss of neutral diversity at linked sites. If hitchhiking affects the same genomic region repeatedly, the process is referred to as recurrent hitchhiking.
Linked selection: The process by which neutral genetic diversity in the genome is lost as an effect of selection at a linked site. Both background selection and hitchhiking contribute to linked selection.

## Acknowledgements

I am grateful to Stuart J. E. Baird, Alexander Suh, Holger Schielzeth and three reviewers for valuable comments on previous versions of the manuscript.

## References

Arbeithuber, B., Betancourt, A.J., Ebner, T. & Tiemann-Boege, I. (2015). Crossovers are associated with mutation and biased gene conversion at recombination hotspots. Proc Natl Acad Sci USA 112:2109–2114.

Bank, C., Ewing, G.B., Ferrer-Admettla, A., Foll, M. & Jensen, J.D. (2014). Thinking too positive? Revisiting current methods of population genetic selection inference. Trends Genet 30:540–546.

Baudat, F., Buard, J., Grey, C., Fledel-Alon, A., Ober, C., Przeworski, M., et al. (2010). PRDM9 Is a Major Determinant of Meiotic Recombination Hotspots in Humans and Mice. Science 327:836–840.

Berner, D. & Salzburger, W. (2015). The genomics of organismal diversification illuminated by adaptive radiations. Trends Genet 31:491–499.

Bhatia, G., Patterson, N., Sankararaman, S. & Price, A.L. (2013). Estimating and interpreting FST: The impact of rare variants. Genome Res 23:1514–1521.

Buckleton, J., Curran, J., Goudet, J., Taylor, D., Thiery, A. & Weir, B.S. (2016). Population-specific FST values for forensic STR markers: A worldwide survey. Forensic Science International: Genetics 23:91–100.

Burri, R. in press. Dissecting differentiation landscapes: A linked selection’s perspective. J Evol Biol.

Burri, R. in review. Linked selection, demography, and the evolution of correlated genomic landscapes in birds and beyond. Mol Ecol.

Burri, R., Nater, A., Kawakami, T., Mugal, C.F., Olason, P.I., Smeds, L., et al. (2015). Linked selection and recombination rate variation drive the evolution of the genomic landscape of differentiation across the speciation continuum of *Ficedula* flycatchers. Genome Res 25:1656–1665.

Charlesworth, B. (2013). Background selection 20 years on. J Hered 104:161–171.

Charlesworth, B. (1998). Measures of divergence between populations and the effect of forces that reduce variability. Mol Biol Evol 15:538–543.

Charlesworth, B., Morgan, M.T. & Charlesworth, D. (1993). The effect of deleterious mutations on neutral molecular variation. Genetics 134:1289–1303.

Comeron, J.M. (2014). Background Selection as Baseline for Nucleotide Variation across the*Drosophila* Genome. PLoS Genet 10:e1004434.

Corbett-Detig, R.B., Hartl, D.L. & Sackton, T.B. (2015). Natural Selection Constrains Neutral Diversity across A Wide Range of Species. PLoS Biol 13:e1002112.

Cruickshank, T.E. & Hahn, M.W. (2014). Reanalysis suggests that genomic islands of speciation are due to reduced diversity, not reduced gene flow. Mol Ecol 23:3133–3157.

Cutter, A.D. & Payseur, B.A. (2013). Genomic signatures of selection at linked sites: unifying the disparity among species. Nat Rev Genet 14:262–274.

Dutoit, L., Vijay, N., Mugal, C.F., Bossu, C.M., Burri, R., Wolf, J.B.W., et al. (2017). Covariation in levels of nucleotide diversity in homologous regions of the avian genome long after completion of lineage sorting. Proc Roy Soc B 284.

Ellegren, H. & Galtier, N. (2016). Determinants of genetic diversity. Nat Rev Genet 17:422–433.

Elyashiv, E., Sattath, S., Hu, T.T., Strutsovsky, A., McVicker, G., Andolfatto, P., et al. (2016). A Genomic Map of the Effects of Linked Selection in Drosophila. PLoS Genet 12:e1006130.

Eyre-Walker, A. & Keightley, P.D. (2007). The distribution of fitness effects of new mutations. Nat Rev Genet 8:610–618.

Feder, J.L., Xie, X., Rull, J., Velez, S., Forbes, A., Leung, B., et al. (2005). Mayr, Dobzhansky, and Bush and the complexities of sympatric speciation in Rhagoletis. Proc Natl Acad Sci USA 102:6573–6580.

Ferrer-Admetlla, A., Liang, M., Korneliussen, T. & Nielsen, R. (2014). On Detecting Incomplete Soft or Hard Selective Sweeps Using Haplotype Structure. Mol Biol Evol 31:1275–1291.

Feulner, P.G.D., Chain, F.J.J., Panchal, M., Huang, Y., Eizaguirre, C., Kalbe, M., et al. (2015). Genomics of Divergence along a Continuum of Parapatric Population Differentiation. PLoS Genet 11:e1004966.

Flowers, J.M., Molina, J., Rubinstein, S., Huang, P., Schaal, B.A. & Purugganan, M.D. (2012). Natural Selection in Gene-Dense Regions Shapes the Genomic Pattern of Polymorphism in Wild and Domesticated Rice. Mol Biol Evol 29:675–687.

Francioli, L.C., Polak, P.P., Koren, A., Menelaou, A., Chun, S., Renkens, I., et al. (2015). Genome-wide patterns and properties of de novo mutations in humans. Nat Genet 47:822–826.

Fraser, B.A., Künstner, A., Reznick, D.N., Dreyer, C. & Weigel, D. (2015). Population genomics of natural and experimental populations of guppies (Poecilia reticulata). Mol Ecol 24:389–408.

Haasl, R.J. & Payseur, B.A. (2016). Fifteen years of genomewide scans for selection: trends, lessons and unaddressed genetic sources of complication. Mol Ecol 25:5–23.

Hey, J. (1991). The structure of genealogies and the distribution of fixed differences between DNA sequence samples from natural populations. Genetics 128:831–840.

Hodgkinson, A. & Eyre-Walker, A. (2011). Variation in the mutation rate across mammalian genomes. Nat Rev Genet 12:756–766.

Huber, C.D., DeGiorgio, M., Hellmann, I. & Nielsen, R. (2016). Detecting recent selective sweeps while controlling for mutation rate and background selection. Mol Ecol 25:142–156.

Hudson, R.R., Slatkin, M. & Maddison, W.P. (1992). Estimation of Levels of Gene Flow From DNA Sequence Data. Genetics 132:583–589.

Irwin, D.E., Alcaide, M., Delmore, K.E., Irwin, J.H. & Owens, G.L. (2016). Recurrent selection explains parallel evolution of genomic regions of high relative but low absolute differentiation in a ring species. Mol Ecol 25:4488–4507.

Jones, F.C., Grabherr, M.G., Chan, Y.F., Russell, P., Mauceli, E., Johnson, J., et al. (2012). The genomic basis of adaptive evolution in threespine sticklebacks. Nature 484:55–61.

Kaplan, N., Hudson, R. & Langley, C. (1989). The hitchhiking effect revisited. Genetics 123:887 – 899.

Kawakami, T., Smeds, L., Backström, N., Husby, A., Qvarnström, A., Mugal, C.F., et al. (2014). A high-density linkage map enables a second-generation collared flycatcher genome assembly and reveals the patterns of avian recombination rate variation and chromosomal evolution. Mol Ecol 23:4035–4058.

Kronforst, Marcus R., Hansen, Matthew E.B., Crawford, Nicholas G., Gallant, Jason R., Zhang, W., Kulathinal, Rob J., et al. (2013). Hybridization Reveals the Evolving Genomic Architecture of Speciation. Cell Reports 5:666–677.

Lohmueller, K.E., Albrechtsen, A., Li, Y., Kim, S.Y., Korneliussen, T., Vinckenbosch, N., et al. (2011). Natural Selection Affects Multiple Aspects of Genetic Variation at Putatively Neutral Sites across the Human Genome. PLoS Genet 7:e1002326.

Martin, S.H., Dasmahapatra, K.K., Nadeau, N.J., Salazar, C., Walters, J.R., Simpson, F., et al. (2013). Genome-wide evidence for speciation with gene flow in Heliconius butterflies. Genome Res 23:1817–1828.

Maynard Smith, J. & Haigh, J. (1974). The hitch-hiking effect of a favourable gene. Genet Res 23:23–35.

McVicker, G., Gordon, D., Davis, C. & Green, P. (2009). Widespread Genomic Signatures of Natural Selection in Hominid Evolution. PLoS Genet 5:e1000471.

Munch, K., Nam, K., Schierup, M.H. & Mailund, T. (2016). Selective sweeps across twenty millions years of primate evolution. Mol Biol Evol 33:3065–3074.

Nachman, M.W. & Payseur, B.A. (2012). Recombination rate variation and speciation: theoretical predictions and empirical results from rabbits and mice. Phil Trans R Soc B 367:409–421.

Nam, K. & Ellegren, H. (2012). Recombination Drives Vertebrate Genome Contraction. PLoS Genet 8:e1002680.

Nei, M. (1972). Genetic Distance between Populations. Am Nat 106:283–292.

Nei, M. & Li, W.-H. (1979). Mathematical model for studying genetic variation in terms of restriction endonucleases. Proc Natl Acad Sci USA 79:5269–5273.

Noor, M.A.F. & Bennett, S.M. (2009). Islands of speciation or mirages in the desert[quest] Examining the role of restricted recombination in maintaining species. Heredity 103:439–444.

Ohta, T. (2002). Near-neutrality in evolution of genes and gene regulation. Proc Natl Acad Sci USA 99:16134–16137.

Oliver, P.L., Goodstadt, L., Bayes, J.J., Birtle, Z., Roach, K.C., Phadnis, N., et al. (2009). Accelerated Evolution of the Prdm9 Speciation Gene across Diverse Metazoan Taxa. PLoS Genet 5:e1000753.

Payseur, B.A. & Nachman, M.W. (2002). Gene density and human nucleotide polymorphism. Mol Biol Evol 19:336–340.

Pennisi, E. (2014). Disputed islands. Science 345:611–613.

Phung, T.N., Huber, C.D. & Lohmueller, K.E. (2016). Determining the Effect of Natural Selection on Linked Neutral Divergence across Species. PLoS Genet 12:e1006199.

Renaut, S., Grassa, C.J., Yeaman, S., Moyers, B.T., Lai, Z., Kane, N.C., et al. (2013). Genomic islands of divergence are not affected by geography of speciation in sunflowers. Nat Commun 4:1827.

Renaut, S., Owens, G.L. & Rieseberg, L.H. (2014). Shared selective pressure and local genomic landscape lead to repeatable patterns of genomic divergence in sunflowers. Mol Ecol 23:311–324.

Roesti, M., Hendry, A.P., Salzburger, W. & Berner, D. (2012). Genome divergence during evolutionary diversification as revealed in replicate lake–stream stickleback population pairs. Mol Ecol 21:2852–2862.

Roesti, M., Kueng, B., Moser, D. & Berner, D. (2015). The genomics of ecological vicariance in threespine stickleback fish. Nat Commun 6:8767.

Roesti, M., Moser, D. & Berner, D. (2013). Recombination in the threespine stickleback genome—patterns and consequences. Mol Ecol 22:3014–3027.

Sabeti, P.C., Varilly, P., Fry, B., Lohmueller, J., Hostetter, E., Cotsapas, C., et al. (2007). Genome-wide detection and characterization of positive selection in human populations. Nature 449:913–918.

Schlötterer, C. (2015). Genes from scratch - the evolutionary fate of *de novo* genes. Trends Genet 31:215–219.

Shriver, M., Kennedy, G., Parra, E., Lawson, H., Sonpar, V., Huang, J., et al. (2004). The genomic distribution of population substructure in four populations using 8,525 autosomal SNPs. Hum Genomics 1:274 – 286.

Singhal, S., Leffler, E.M., Sannareddy, K., Turner, I., Venn, O., Hooper, D.M., et al. (2015). Stable recombination hotspots in birds. Science 350:928–932.

Smukowski, C.S. & Noor, M.A.F. (2011). Recombination rate variation in closely related species. Heredity.

Soria-Carrasco, V., Gompert, Z., Comeault, A.A., Farkas, T.E., Parchman, T.L., Johnston, J.S., et al. (2014). Stick Insect Genomes Reveal Natural Selection’s Role in Parallel Speciation. Science 344:738–742.

Stephan, W. (2010). Genetic hitchhiking versus background selection: the controversy and its implications. Phil Trans R Soc B 365:1245–1253.

Tang, K., Thornton, K.R. & Stoneking, M. (2007). A New Approach for Using Genome Scans to Detect Recent Positive Selection in the Human Genome. PLoS Biol 5:e171.

Terekhanova, N.V., Seplyarskiy, V.B., Soldatov, R.A. & Bazykin, G.A. (2017). Evolution of Local Mutation Rate and Its Determinants. Mol Biol Evol.

Turner, T.L. & Hahn, M.W. (2010). Genomic islands of speciation or genomic islands and speciation? J Mol Ecol 19:848–850.

Turner, T.L., Hahn, M.W. & Nuzhdin, S.V. (2005). Genomic Islands of Speciation in *Anopheles gambiae*. PLoS Biol 3:e285.

Van Doren, B.M., Campagna, L., Helm, B., Illera, J.C., Lovette, I.J. & Liedvogel, M. (2017). Correlated patterns of genetic diversity and differentiation across an avian family. Mol Ecol 26:XXX–XXX.

Vijay, N., Bossu, C.M., Poelstra, J.W., Weissensteiner, M.H., Suh, A., Kryukov, A.P., et al. (2016). Evolution of heterogeneous genome differentiation across multiple contact zones in a crow species complex. Nat Commun 7:13195.

Vijay, N., Weissensteiner, M., Burri, R., Kawakami, T., Ellegren, H. & Wolf, J.B.W. (2017). Genome-wide signatures of genetic variation within and between populations - a comparative perspective. bioRxiv.

Voight, B.F., Kudaravalli, S., Wen, X. & Pritchard, J.K. (2006). A Map of Recent Positive Selection in the Human Genome. PLoS Biol 4:e72.

Weir, B.S. & Cockerham, C.C. (1984). Estimating F-Statistics for the Analysis of Population-Structure. Evolution 38:1358–1370.

Whitlock, M.C. (2011). G’ST and D do not replace FST. Mol Ecol 20:1083–1091.

Yi, X., Liang, Y., Huerta-Sanchez, E., Jin, X., Cuo, Z.X.P., Pool, J.E., et al. (2010). Sequencing of 50 Human Exomes Reveals Adaptation to High Altitude. Science 329:75–78.

Zeng, K. & Charlesworth, B. (2011). The Joint Effects of Background Selection and Genetic Recombination on Local Gene Genealogies. Genetics.

Zeng, K. & Corcoran, P. (2015). The Effects of Background and Interference Selection on Patterns of Genetic Variation in Subdivided Populations. Genetics 201:1539–1554.

